# Robin’s Viewer: Using Deep-Learning Predictions to Assist EEG Annotation

**DOI:** 10.1101/2022.08.07.503090

**Authors:** Robin Weiler, Marina Diachenko, Erika Juarez-Martinez, Arthur-Ervin Avramiea, Peter Bloem, Klaus Linkenkaer-Hansen

## Abstract

Machine learning techniques such as deep learning have been increasingly used to assist EEG annotation, by automating artifact recognition, sleep staging, and seizure detection. In lack of automation, the annotation process is prone to bias, even for trained annotators. On the other hand, completely automated processes do not offer the users the opportunity to inspect the models’ output and re-evaluate potential false predictions. As a first step towards addressing these challenges, we developed Robin’s Viewer (RV), a Python-based EEG viewer for annotating time-series EEG data. The key feature distinguishing RV from existing EEG viewers is the visualization of output predictions of deep-learning models trained to recognize patterns in EEG data. RV was developed on top of the plotting library Plotly, the app-building framework Dash, and the popular M/EEG analysis toolbox MNE. It is an open-source, platform-independent, interactive web application, which supports common EEG-file formats to facilitate easy integration with other EEG toolboxes. RV includes common features of other EEG viewers, e.g., a view-slider, tools for marking bad channels and transient artifacts, and customizable preprocessing. Altogether, RV is an EEG viewer that combines the predictive power of deep-learning models and the knowledge of scientists and clinicians to optimize EEG annotation. With the training of new deep-learning models, RV could be developed to detect clinical patterns other than artifacts, for example sleep stages and EEG abnormalities.

## 1 Introduction

Electroencephalography (EEG) remains one of the most widely used methods for measuring normal brain activity (Biasiucci et al., 2019; da Silva, 2013; Niedermeyer and da Silva, 2005), essential in the study of brain disorders (e.g., epilepsy, sleep-, and mental disorders (Acharya et al., 2013; Arns et al., 2013; Chen and Koubeissi, 2019; Diaz et al., 2016; Olbrich and Brunovski, 2021)), and more recently, a cornerstone for the development of brain-computer interfaces (BCI) which interpret EEG to establish control over external devices, e.g., prosthetic limbs (Abiri et al., 2019; Allison et al., 2007; Wolpaw et al., 1991). EEG signals, however, come mixed with many physiological and non-physiological artifacts (e.g., muscle activity, eye movements and blinking for the former, and electrode movement, impedance changes, and interference from other electronic devices) (Barua and Begum, 2014; Niedermeyer and da Silva, 2005). The frequency spectra of artifacts may overlap with those of neural oscillations relevant for clinical evaluation (e.g., epilepsy) or scientific research (Donoghue et al., 2022; McKay and Tatum, 2019; Ward, 2003). Therefore, accurate artifact detection and removal is crucial prior to any qualitative or quantitative EEG assessment. Generally, this requires the input of experienced clinicians and/or researchers, which can be time consuming and, inevitably, introduces a certain degree of subjectivity (Biasiucci et al., 2019).

Machine-learning techniques trained to detect artifacts in EEG data can address the aforementioned issues, by providing an objective standard for artifact marking, and speeding up the marking process. The most common machine-leaning techniques used for artifact detection in EEG data, are based on support vector machines (SVMs) (Barua and Begum, 2014; Sai et al., 2017; Shao et al. 2008), k-nearest neighbor classifiers (k-NN) (Barua and Begum, 2014; Roy, 2019), independent component analysis (ICA) (Barua and Begum, 2014; Radüntz et al., 2015; Sai et al., 2017), and, as of recently, various deep-learning models (e.g., autoencoders (Roy et al., 2019; Yang et al., 2016; 2018), convolutional neural networks (CNNs) (Diachenko et al., 2022; Jurczak et al., 2022; Roy et al., 2019; Sun et al., 2020), and recurrent neural networks (RNNs) (Liu et al., 2022; Roy et al., 2019)). Deep-learning solutions, in particular, have increased in popularity for artifact handling (Craig et al., 2019; Roy et al., 2019) due to the minimal preprocessing they require and because of their ability to learn very complex functions between input data and the desired output classification (LeCun et al., 2015). Nevertheless, the performance of automated EEG annotation through deep-learning models is still far from replacing trained human annotators in a clinical setting. Furthermore, when artifact removal is completely automated, it does not offer the users the opportunity to inspect the models’ output and re-evaluate potential false predictions.

To this end, we developed Robin’s Viewer (RV), a user-friendly EEG viewer for annotating time-series EEG data using deep-learning models trained to recognize artifacts in EEG data, as a decision-support-system. As such, with RV, we combine the predictive power of deep-learning models and the knowledge of subject-matter experts (i.e., scientists and clinicians) to improve the EEG annotation process. Importantly, RV was developed in close collaboration with machine-learning experts, neurologists, EEG researchers, and non-expert users (e.g., students), to ensure comfortable and efficient EEG viewing and annotation. RV is open-source (MIT license), platform independent (it runs in the user’s web browser), and supports many common EEG-file formats, allowing for integration into data-processing pipelines developed for other EEG toolboxes. RV was developed in Python and builds on the popular M/EEG analysis toolbox MNE (Gramfort et al., 2013), Plotly^1^, a Python-based plotting library, and Dash^2^, used for the interactive graphical user interface (GUI) and for providing a locally hosted server to run the application. In RV, the user can load EEG recordings, apply common preprocessing methods (e.g., bandpass filtering, re-referencing, and down-sampling), plot the data, use the GUI to navigate through the signal, and mark bad channels and artifacts. To provide an example of how the viewer can be used with integrated deep-learning predictions, we have additionally included with RV a proof-of-concept deep-learning model trained to recognize artifacts in resting-state, multichannel EEG recordings (Diachenko et al., 2022). RV can be downloaded for free at https://github.com/RobinWeiler/RV.git, where we also provide code documentation, demos, and example data. Here, we give an overview of the design and key features of RV using artifact annotation as the running-example.

## 2 Methods

### 2.1 Programming Environment

We developed RV in Python and built it around the following main libraries: Plotly, Dash, and MNE. A full list of the required libraries can be found in the “requirements.txt” file included on GitHub. Because of Python’s portability and the fact that Plotly and Dash allow RV to be served in the user’s web browser, our application is largely platform independent. We tested RV on the three major platforms MacOS, Windows, and (Ubuntu) Linux, as well as the popular web browsers Safari, Google Chrome, and Mozilla Firefox.

All plots and visualizations, as well as parts of the interactive GUI, including the slider and artifact annotation mechanism, included in RV, were written using Plotly. We used Dash to deploy and host the application in the user’s web browser. Moreover, most of the buttons, dropdown-menus, and pop-up windows of RV were developed in Dash using the integrated callbacks.

MNE (Gramfort et al., 2013) is a popular M/EEG analysis toolbox built in Python, which features a wide variety of common techniques applied to EEG data. RV relies on the data structures provided by MNE. Additionally, all the preprocessing applied in RV depends on MNE code, including automatic bad-channel detection, which uses the Autoreject library (Jas et al., 2017), and which is also based on MNE. However, when needed, the user can pre-process their data outside of RV and then plot it directly using the techniques outlined in Section 3.8, skipping the preprocessing steps applied by RV.

### 2.2 Deep-Learning Model

A convolutional neural network was trained on expert-annotated resting-state EEG data from typically developing children and children with neurodevelopmental disorders, with the purpose of recognizing artifacts (Diachenko et al., 2022). This model was integrated into RV as a proof-of-concept decision-support system for EEG artifact annotation. The model reached 80% balanced accuracy (a performance metric based on the average of sensitivity and specificity which is used to assess accuracy in imbalanced data, i.e., where one target class appears more often than the other) in distinguishing artifact from non-artifact EEG patterns and, importantly, detected valid artifacts missed by the trained annotator. Therefore, using this model together with RV could increase the quality and speed of manual annotation of EEG artifacts.

RV also allows the model to be used for automatic artifact annotation, i.e., it can automatically annotate portions of the signal predicted with confidence that exceeds a customizable threshold, which will be explained in more detail in Section 3.5. For the data where the model confidence does not cross the fixed threshold, the user has to make the final decision.

Importantly, the model applies the following preprocessing steps, independent of the preprocessing set by the user in RV, to generate its predictions: First, the data is bandpass filtered in the range of 0.5 Hz to 45 Hz with a Hamming window. Then, bad channels, which should either be marked through RV (as explained in Section 3.4) or be contained in the loaded data, are interpolated. Finally, 19 channels are selected according to the standard 10-20 system and the data is segmented into 1-second segments with 0.5 seconds overlap. From these segments, time-frequency plots are generated which the model uses as input. For more details, we refer to Diachenko et al. (2022).

## 3 Results

In the following, we give an overview of the design and key features of RV and its graphical user interface (GUI) using artifact annotation as the running-example.

### 3.1 Loading EEG Data

The initial screen of RV consists of a pop-up window where the user can load EEG data and select from the preprocessing and visualization methods provided by the application. This window can be seen in Figure 1 and can always be reopened with the “Preprocessing” button in the top-left corner of the menu bar of the main GUI (seen in Figure 2A and explained in Section 3.3.1).

**Figure 1.**
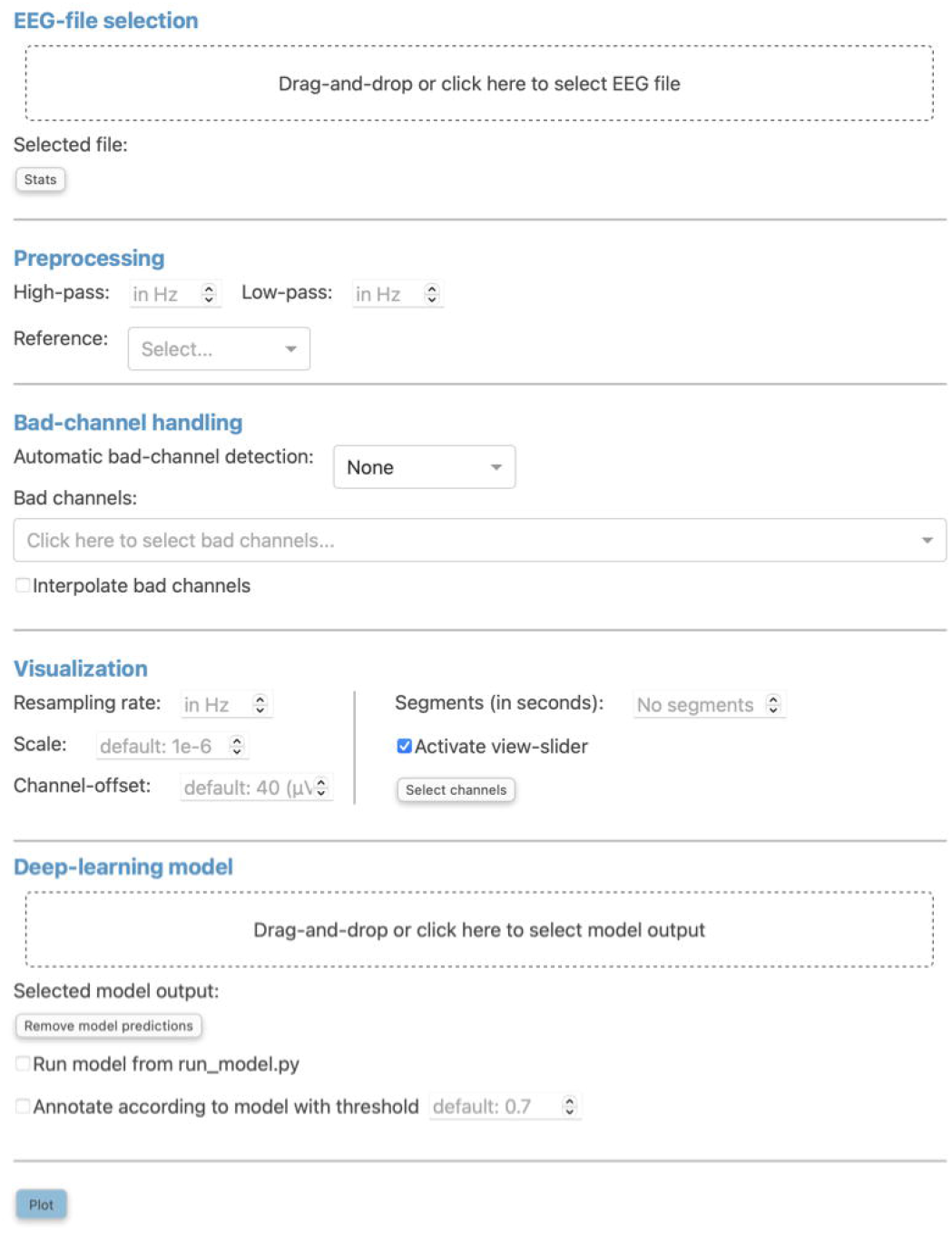
The “Preprocessing” pop-up window is the initial screen of RV and has five distinct sections: First, the EEG-file selection (see Section 3.1), where the user can load their EEG recording and display selected statistics about the data once it is loaded. Second, the preprocessing settings (see Section 3.2), which are used to set a bandpass filter (FIR filter with Blackman window) and a custom reference. Third, the bad-channel handling (see Section 3.2 and 3.4), where the user can decide whether to use automatic bad channel detection, and whether to interpolate bad channels. Fourth, the visualization settings (see Section 3.2), comprised of downsampling, custom scaling (by default 1e-6 as RV scales data from volts to microvolts for plotting), the gap between traces (by default 40 (μV); setting this to 0 results in butterfly mode where all traces are collapsed on top of each other; values higher than 40 move traces further apart), segment length to plot (by default 60 (seconds)), whether or not to activate the view-slider, and selection of channels to plot. Visualization settings will only be applied to the data for plotting and hence will not be saved in the save-file (in contrast to the preprocessing settings). Fifth, the deep-learning model settings (see Sections 3.2 and 3.5), where previously saved model output can be loaded and the integrated deep-learning model can be activated to generate predictions. Clicking the “Plot” button at the bottom will close this window and, after a loading screen (which lasts as long as it takes to plot the data), it will open up the main GUI.

**Figure 2.**
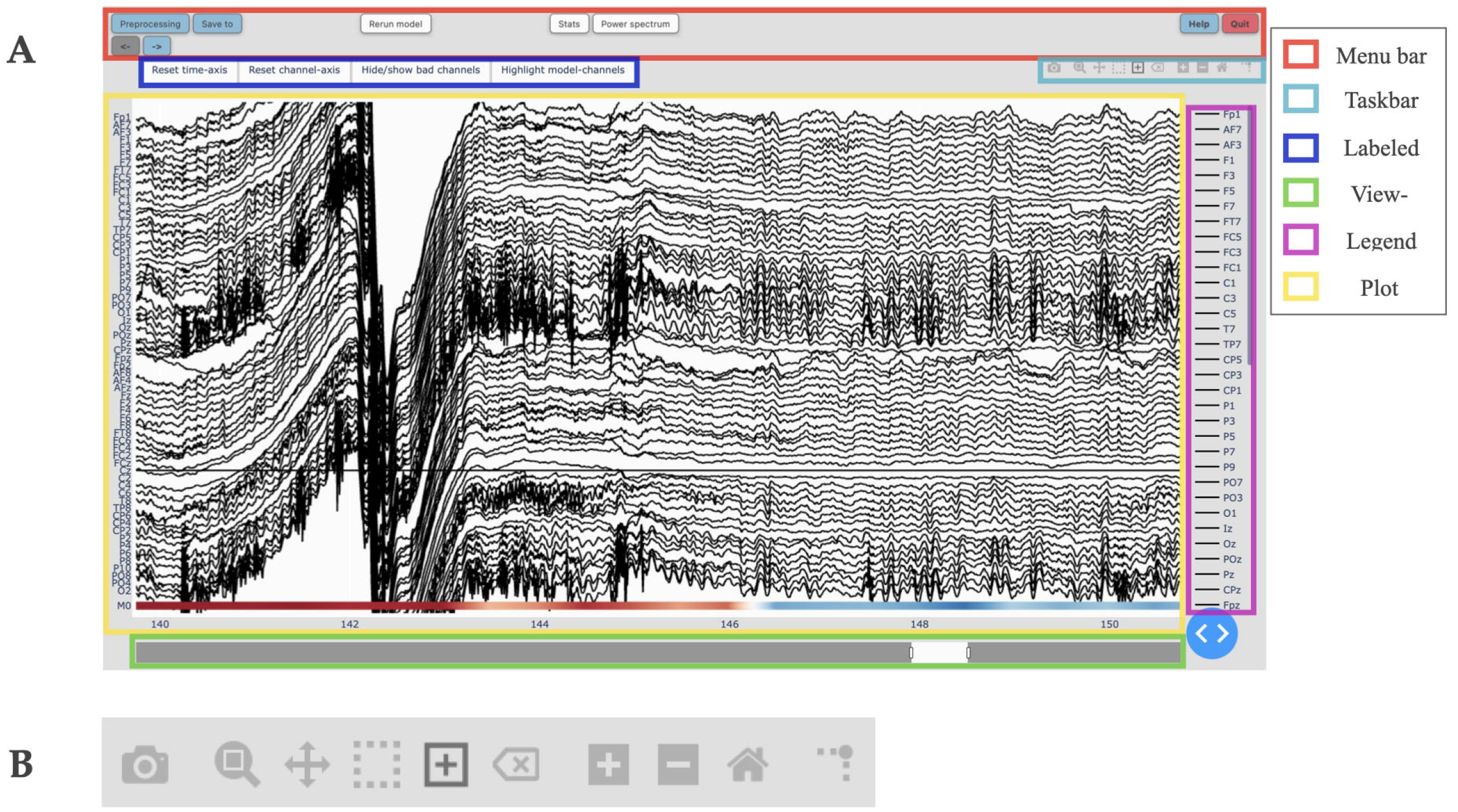
(A) The main Graphical User Interface (GUI) of RV with its six subsections highlighted (see Section 3.3): The menu bar (*red*) contains buttons to open pop-up windows for file loading and preprocessing, saving, annotation statistics, power spectrum, help, and shutting down RV, as well as a button to recalculate the deep-learning model’s predictions (if activated) and left and right arrow buttons used to navigate the currently viewed timeframe (if the data is loaded in segments). The taskbar (*light blue*) seen in detail in B. The labeled buttons (*dark blue*): The first two buttons are used to reset the view, either across the channel- (“Reset channel-axis”) or time-axis (“Reset time-axis”). The third button (“Hide/show bad channels”) allows to hide marked bad channels from the plot. The fourth button (“Highlight model-channels”) only appears when deep-learning predictions are activated and highlights the channels used by the model to make its predictions in blue. The view-slider (*green*) is used to scroll continuously along the time-axis of the signal. The legend (*purple*) shows all available channels. Clicking on any channel in the legend once will hide it from the plot (clicking again reverses this). Double-clicking on a channel will hide all other channels from the plot, except for the selected channel. The plot (*yellow*) shows the traces for all selected channels spread across the vertical axis, and time (in seconds) on the horizontal axis. The user can hover over any given point in the plot in order to display the time (in seconds) and amplitude (in μV if no custom scaling was used) values of the trace under the mouse. The deep-learning predictions, if activated (see Section 3.5), are plotted below the EEG traces. (B) The taskbar hosts ten buttons from left to right: 1. Take a picture of the (annotated) EEG signal and download it as a .png file; 2. Select an area to zoom in on; 3. Move view along the channel- or time-axis; 4. Select a segment of the data for which to calculate and display the main underlying frequency and power spectrum; 5. Select a segment of the plot to annotate; 6. Delete the currently selected annotation; 7. Zoom in one step; 8. Zoom out one step; 9. Zoom out as much as necessary to show all channels for the entire duration of the recording (or segment); 10. Display a ruler from both axes to the datapoint currently hovered on.

RV follows the following file structure: There is the “data” directory, from which files can be loaded but not overwritten. This is done to ensure that the user never accidentally overwrites the original raw EEG-data files. Next, there is the “save_files” directory, where all files annotated in RV will be saved, and where files can be overwritten when the user explicitly chooses to do so in the “Save to” pop-up window (more information on RV’s saving functionality in Section 3.7). Additionally, there is the “temp_raw.fif” file located in the main RV directory, which is a temporary working save-file updated automatically with every annotation made to the data and which can be loaded in case RV is closed accidentally without saving.

All files located in both the “data” and “save_files” directory, as well as the “temp_raw.fif” file, can be loaded in RV. Currently, the following file-formats are supported for loading: .edf, .bdf, .fif, and .set. To load these files, the user can either drag-and-drop them into the “Drag-and-drop or click here to select EEG file” area (Figure 1), or click on said area, after which a file-selection window will open and allow the user to select the file to load. However, RV only has access for reading files in the directories named above because the browser RV runs in hides file paths for security reasons. Therefore, the selected file will still have to be inside one of the predefined directories to be loadable. Note that the user can also choose to load the data outside of RV (e.g., to load a file-format not currently supported) and pass it as a MNE Raw data-structure to the “run_viewer” function (explained in more detail in Section 3.8). Using this function, the user can circumvent RV’s restriction of only loading files from the predefined directories, and only saving files to the “save_files” directory, by loading the data externally and passing it to the “run_viewer” function, together with a desired save-file path.

After loading EEG data, the user can use the “Stats” button (Figure 1) in order to view statistics surrounding the annotated data for a quick overview of the data before plotting (Figure 3). Currently, the implemented statistics consist of the recording’s total length (in seconds), the amount of artifacts and clean data (in seconds), as well as the amount of consecutive, clean intervals longer than two seconds (this is useful because, if most of the remaining clean data segments are shorter than two seconds, the recording’s quality is likely not sufficient for further analysis). Furthermore, we included a histogram to show the distribution of the length of clean data segments. These statistics are intended to give the annotator a feeling for the quality of the data and whether this data can be used for further analysis. We invite researchers to implement new statistics, or to create a feature request on GitHub.

**Figure 3.**
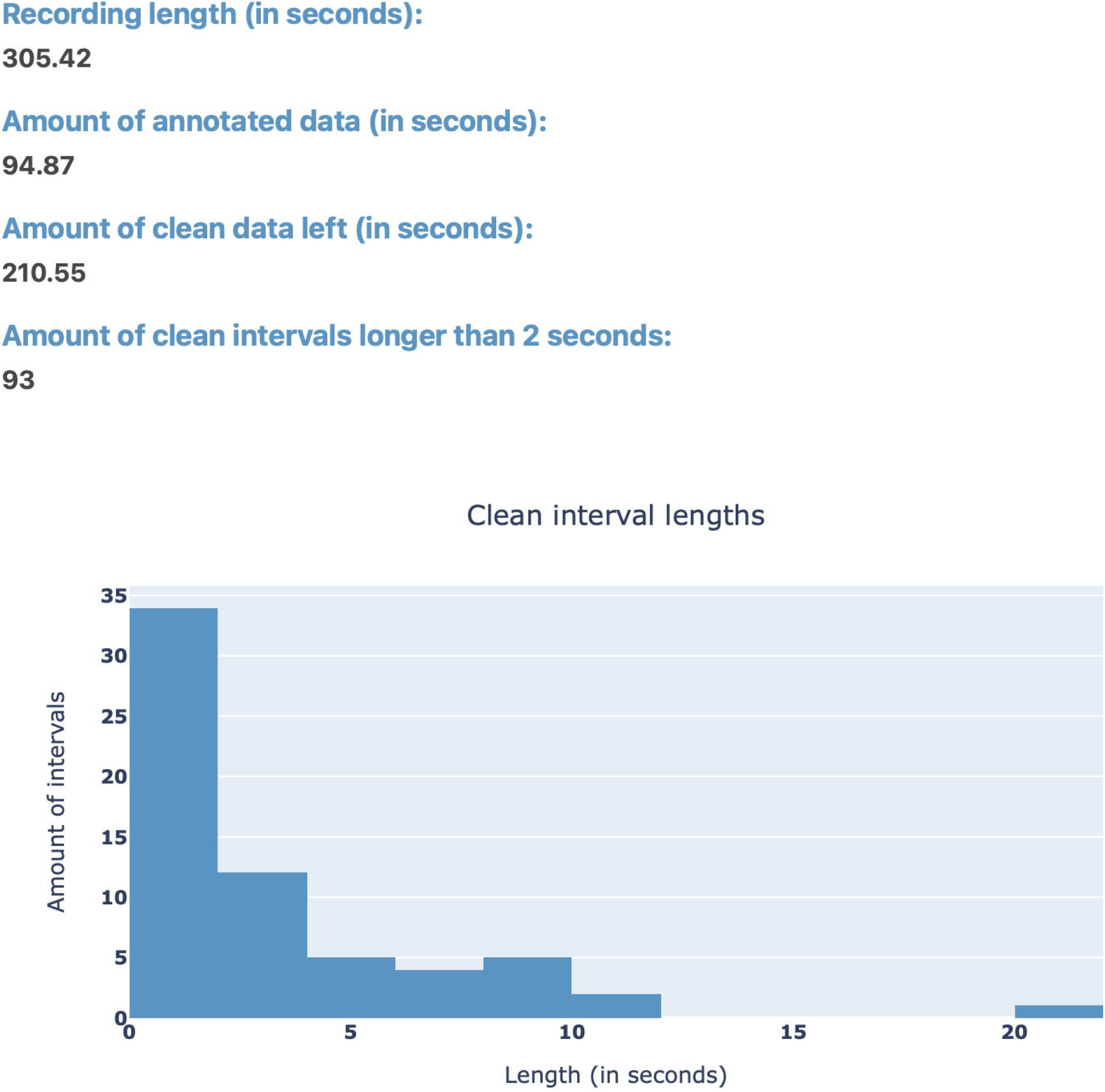
The “Stats” pop-up window offers an overview over the annotated data. Currently, the integrated statistics consist of the recording’s total length (in seconds), the amount of annotated (artifact) and non-annotated (artifact-free) data (in seconds), as well as the amount of consecutive, non-annotated (artifact-free) intervals longer than two seconds. Finally, there is a histogram to show the distribution of the length of non-annotated data.

### 3.2 Preprocessing and Visualization Settings

RV offers a handful of popular EEG preprocessing methods within the “Preprocessing” pop-up window seen in Figure 1, which rely on the MNE package (Gramfort et al., 2013). These include bandpass filtering, re-referencing, bad channel interpolation, and automatic bad-channel detection, which is based on the RANSAC method of the PREP pipeline (Bigdely-Shamlo et al., 2015) implemented in the Autoreject library (Jas et al., 2017). The flowchart in Figure 4 shows the order in which preprocessing and visualization settings are applied to the data.

**Figure 4.**
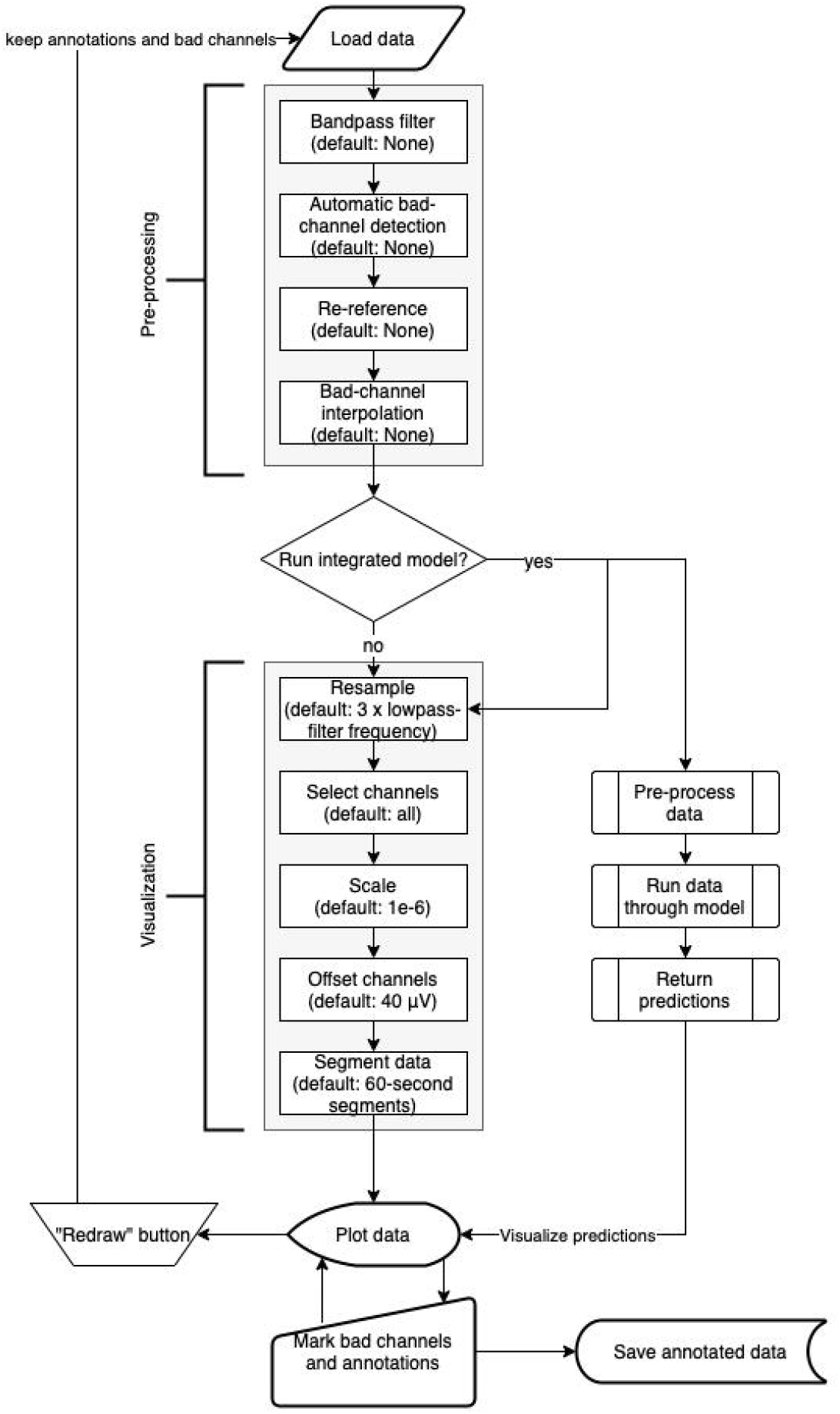
Flowchart of RV’s preprocessing and visualization settings pipeline (see Section 3.2). After the data is loaded, it goes through the preprocessing and visualization steps in the following order before being plotted: Bandpass-filtering (FIR filter with Blackman window) (none by default), automatic bad channel detection (RANSAC method of the PREP pipeline (Bigdely-Shamlo et al., 2015) implemented in the Autoreject library (Jas et al., 2017)) (none by default), re-referencing (none by default), bad-channel interpolation (none by default), resampling (three times the lowpass-filter frequency (if given) by default), channel selection (all by default), scaling (1e-6 by default), channel offsetting (40 μV by default), and data segmentation (60-second segments by default). Importantly, many of these steps are optional (as indicated by their default values) and will be skipped if they are not specifically set. In parallel to the visualization settings, the deep-learning model generates predictions (if activated). Using the “Redraw” button in the menu bar of the main GUI (see Figure 2A) restarts the pipeline but keeps annotations and marked bad channels. Saving the annotated data will include the preprocessing settings but not the visualization settings.

For the high- and low-pass filter, the user needs to provide the values (in Hz) in the respective input fields. For this bandpass filter, we used the “mne.io.Raw.filter” function from MNE set to an FIR filter with a Blackman window (Lai, 2003) (all the other parameters were MNE defaults). More filters will be added in future versions of RV. If the loaded EEG data has previously been preprocessed (because it is a save-file or because it was preprocessed outside of RV), the filter settings will be extracted and filled in automatically. In this case, if no changes are made to these values, or fields are empty, the data will be loaded without any additional filtering.

Below the bandpass filter, the user can choose how to re-reference the signals: using an average reference, a Cz-reference, or altogether skipping this step by selecting no custom reference. Importantly, when the annotated data is saved in RV, this includes the applied preprocessing options (bandpass filter and custom reference). This is to avoid the situation where, e.g., high-frequency artifacts are filtered away during viewing but not when subsequently applying quantitative analysis algorithms.

Below the preprocessing options, there is a compartment for bad-channel handling. For automatic bad-channel detection, there are two options: skip the step altogether (“None”) or use automatic bad-channel detection (“RANSAC”). For more details on the automatic bad-channel detection, see Section 3.4.2. More methods for automatic bad-channel detection will be added in future versions of RV. Bad channels can also be marked manually in RV, which will be explained in Section 3.4.1. If the loaded file already contains marked bad channels, they will be selected in the bad-channel dropdown-menu. The user can de-select channels here or select more channels by clicking on the dropdown-menu, which will list all available EEG channels. The user can select a channel or type the beginning of a channel name in the dropdown-menu text box, which will filter the list of available channels.

Interpolation of bad channels can be activated in this compartment as well and is performed when there are marked bad channels (regardless of whether they were marked manually or by the automatic bad-channel detection algorithm) and when the loaded file contains channel-position information. If the latter is not provided, RV will display an error message.

Aside from the preprocessing options and bad-channel handling, there are several settings that only affect the visualization and are not reflected in the save-file when saved. First, the user can determine the resampling frequency (in Hz) used for plotting the signal. By default, this is initialized with three times the lowpass-filter parameter (leaving this field empty will load the signal with its original sampling frequency). We highly recommend downsampling the data before plotting, as this will have a big impact on the overall performance and usability of RV as it effectively reduces the number of datapoints plotted at once. For the future, we will investigate the option to automatically adapt the sampling rate based on the current level of zoom, e.g., by plotting more points as the user zooms in on the data. Note that resampling will only be used for plotting and does not affect the sampling frequency of the data saved in the end.

Another option that affects the visualization of the signal is the scaling factor, which determines the scaling the data should have in relation to volts, the default unit used by MNE. By default, this is equal to 1e-6, allowing for EEG data to be scaled from volts to microvolts so that fluctuations in the signal are visible. Additionally, there is an input field allowing the user to change the space between channels for plotting, called “Channel-offset”. By default, this is equal to 40 μV, which creates enough space between individual channels while still spreading them out as much as possible to preserve detail in the traces. Butterfly mode, i.e., setting channel offset to 0, will collapse all channels on top of each other, which can be useful to identify potential bad channels with high variance. Setting channel offset to numbers bigger than 40 will spread the channels further apart, requiring the user to scroll up and down through the channels.

Another visualization option that significantly improves RV’s performance and responsiveness is the ability to load and visualize the recording in segments of a specified length (in seconds). When using this option, the user can move between the segments (which have half a second overlap in either direction) using the left- and right-arrow buttons in the menu bar of the GUI (Figure 2A), just below the “Preprocessing” button. When the recording is segmented in this way, fewer datapoints are plotted at once, compared to the case where the entire signal is loaded. By default, the recording is segmented into 60-second segments plus half a second overlap with both the previous and following segment. On a MacBook Pro 13-inch, 2020, with the Apple M1 chip and 16GB RAM, for example, switching between two 60-second (plus 1-second overlap) segments with 32 channels and a sample frequency of 135 Hz (i.e. 263,520 points plotted at once) takes approximately one second.

Underneath the input field to set the segment size, there is an option to enable/disable the adjustable view-slider, which can be used to scroll through the recording (or segments) continuously. Using the view-slider, the user could load their EEG recording in segments of 60 seconds, for example, and then continuously scroll through each segment with an adjustable (see Section 3.3.4) sliding window (i.e., without switching between segments until the end of a (e.g., 60-second) segment is reached). Scrolling through the data continuously comes with the cost of data outside of the timeframe shown by the view-slider having to be preloaded in the background (e.g., the full 60-second segment is preloaded even if only ten seconds are displayed at a time). Therefore, disabling the view-slider, and viewing the signal in smaller (e.g., 10-second) segments instead, can improve RV’s performance, if necessary. By default, the view-slider is active and initialized to span a ten-second timeframe.

Finally, by pressing the “Select channels” button, the user can select what channels they want to plot the signals for (Figure 5). Channels can be selected from a sensor-layout plot (if the loaded data-file contains channel-position information) or by choosing specific channel names to be plotted in a dropdown-menu. If no selection is made, all channels will be plotted and visible after loading.

**Figure 5.**
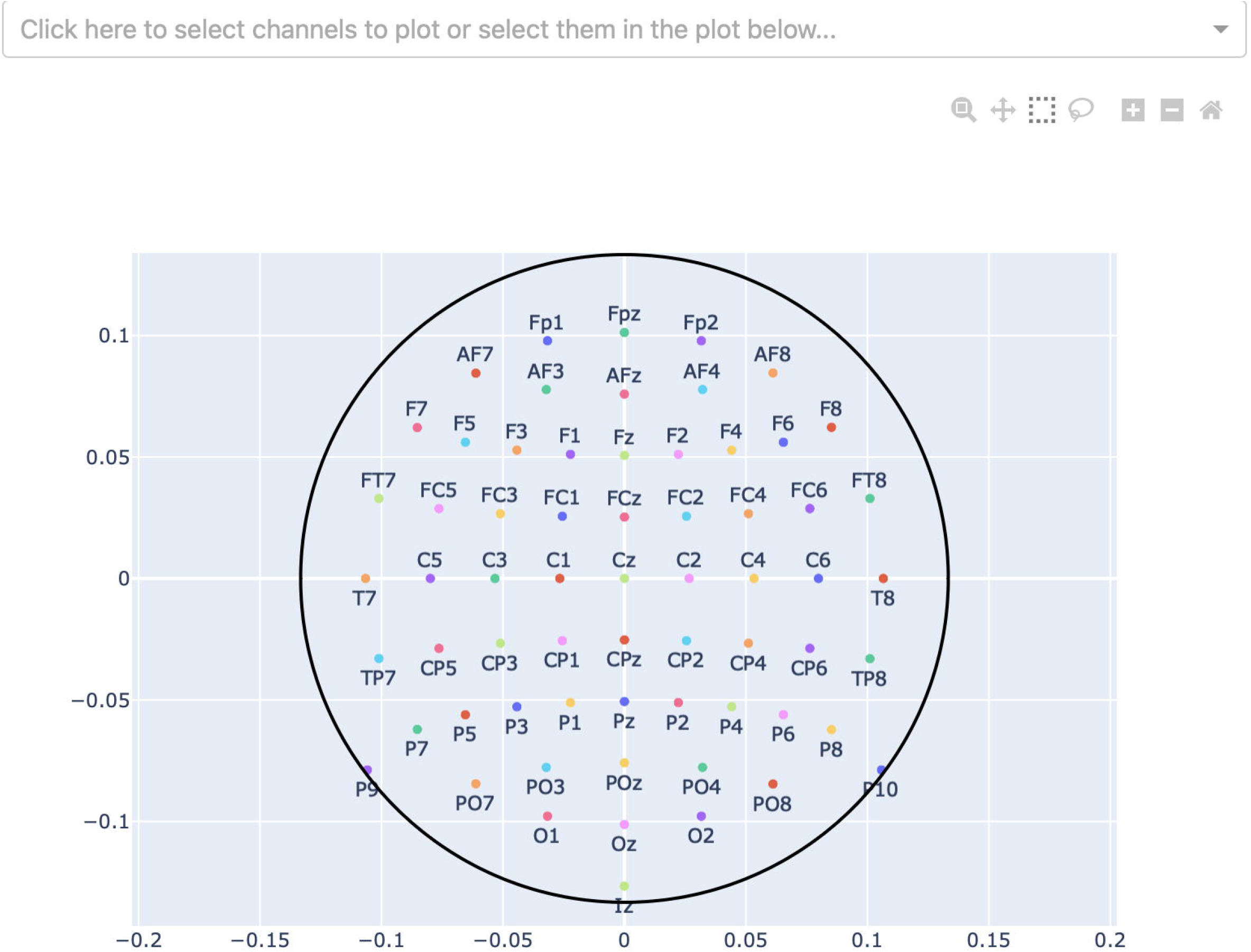
Channel-selection pop-up window where the user can select specific channels to plot. At the top, there is a dropdown-menu to select channels out of a list of all channels. Alternatively, the channels can be selected on the channel layout below by clicking and dragging on the plot.

Pressing the “Plot” button at the bottom will close the “Preprocessing” window and, after a period in which the signal is loaded, it will open the main GUI where the preprocessed EEG data is displayed. The “Preprocessing” window can be reopened at any time using the “Preprocessing” button in the menu bar of the application (Section 3.3.1), and all the preprocessing and visualization settings can be changed. Clicking the “Plot” button again will reset the current view and remove all existing annotations and marked bad channels. This is because changing the bandpass filter settings can influence the timing of artifacts. When needed, all relevant data should first be saved before changing these settings and re-plotting.

### 3.3 Graphical User Interface (GUI)

The GUI of RV is divided into six components, as indicated in Figure 2A, each of which will be explained in detail in the following subsections.

#### 3.3.1 Menu Bar

The menu bar is located at the top of the screen. It contains buttons which, when pressed, open pop-up windows for file loading and preprocessing, file saving, annotation statistics, power spectrum, help, and shutting down RV. Additionally, the menubar hosts a button to rerun the deep-learning model, as well as left and right arrow buttons used to navigate between segments if the data is loaded in segments. The “Preprocessing” button opens the file-selection and preprocessing window mentioned in Section 3.1. The “Save to” button opens a pop-up window allowing the user to save the annotated data with a given filename (explained in detail in Section 3.7).

Next to the “Save to” button, there is the “Re-run model” button, which, as the name suggests, reruns the integrated deep-learning model (see Section 3.5.2), if activated. This button should be used after marking all bad channels in order to recalculate the model’s predictions while considering the newly marked bad channels.

The “Stats” button opens the same pop-up window described in Section 3.1 and shown in Figure 3, containing statistics surrounding the annotated data. Next to the “Stats” button, there is the “Power Spectrum” button, which opens a pop-up window showing the most prominent frequency, as well as the respective power spectrum for the currently selected time interval of EEG data. For the power-spectrum computation, the Welch method (Welch and Peter, 1967) is used as implemented in the Python-library SciPy (Virtanen et al., 2020). How to select such a time interval will be explained in the following Section 3.3.2 - button 4.

Finally, on the top-right, there are two buttons grouped together. On the left, the “Help” button opens a pop-up window containing links to the documentation and GitHub. On the right, the “Quit” button displays a pop-up window asking for confirmation to shut down RV, shuts down the local server RV is running on, and after this the RV interface cannot be used anymore. Before shutting down RV, the user should save the annotated data using the “Save to” button mentioned above (if the user forgets to save the data, it can be restored using the “temp_raw.fif” file after RV is restarted).

#### 3.3.2 Taskbar

The taskbar seen in Figure 2B (located in the upper right corner of the GUI, below the menu bar (Figure 2A)) allows the user to interact with the EEG plot. From left to right, there are the following buttons:

1. Take a picture of the (annotated) EEG signal and download it as a .png file.
2. While this button is active, select an area to zoom in on (click-and-drag).
3. While this button is active, move view along the channel- or time-axis, or both simultaneously (click-and-drag).
4. While this button is active, click on a channel to mark it as a bad channel. After a brief loading period, the respective channel will be marked in gray. Clicking on a bad channel again will remove the marking. Also, while this button is active, select a segment of the data for which to calculate and display the main underlying frequency, as well as a power spectrum in “Power-spectrum” pop-up window, which can be opened using the “Power spectrum” button in the menu bar (click-and-drag). It is possible to select all channels or only a few desired ones, as explained in Section 3.3.5.
5. While this button is active, select a segment of the plot to annotate, which creates a semi-transparent red box (indicating the presence of an artifact) spanning the entire vertical axis in view (currently only annotations across all channels are supported) across the selected time interval (click-and-drag). These annotations can be freely adjusted in their position and size, or removed entirely (with button 6), when clicked on. The annotation is saved with “bad_artifact” as its description (see Section 3.6 for more details). This tool is activated by default.
6. Delete the currently selected annotation.
7. Zoom in one step.
8. Zoom out one step.
9. Zoom out as much as necessary to show all channels for the entire duration of the recording (or segment), including all peaks in the data (potentially produced by artifacts).
10. Display rulers for both axes corresponding the datapoint currently hovered on.

#### 3.3.3 Labeled Buttons

By default, there are three labeled buttons in the top-left of the plot (Figure 2A). The first two buttons are used to reset the view, either across the channel- (“Reset channel-axis”) or time-axis (“Reset time-axis”). “Reset channel-axis” will reset the view to fit all channels in the plot. “Reset time-axis” will reset the plotted timeframe to show the initial configuration, i.e., the first ten seconds of the recording (or segment) if the view-slider is activated or the entire recording (or segment) otherwise.

The third button (“Hide/show bad channels”) allows the user to hide marked bad channels from the plot. Once bad channels have been hidden, they can be made visible by clicking the button again. It is important to note that, after marking bad channels, the user will first have to use the “Redraw” button in the menu bar in order for the changes to have an effect, and for the “Hide/show bad channels” button to work accordingly.

An additional fourth button, called “Highlight model-channels”, appears only when the integrated deep-learning model is activated. This button highlights the channels which were used by the model to generate predictions in blue. These channels are defined in the model implementation and cannot be changed. Knowing which channels are considered by the model can be helpful in understanding its predictions, e.g., when an artifact was not marked by the model because it occurred on a channel that was not part of the model input.

#### 3.3.4 View-Slider

The view-slider, once activated in the “Preprocessing” window (Figure 1), is located at the bottom of the screen (Figure 2A), and can be used to continuously scroll through the recording (horizontally along the time-axis) by clicking and dragging it (as mentioned in Section 3.2). The slider’s range, by default initialized to ten seconds, can be adjusted by clicking and dragging the small rectangles at its edges. In this way, the user can visualize anything from the entire recording (or segment) at once, down to several milliseconds. It should be noted that the same functionality as the view-slider can also be achieved by clicking and dragging on the time-axis’ labels or selecting the third button in the taskbar (Figure 2B – button 3) and then clicking and dragging on the plot directly.

#### 3.3.5 Legend

At the right side of the plot, there is a scrollable legend showing the names of all available EEG channels (Figure 2A). Clicking on any channel name hides the channel from the plot. Double-clicking a channel name hides all channels except for the selected one, and the user can follow up by adding more channels to displayed, by clicking their names once. The latter can be used to retrieve the most prominent underlying frequency and power spectrum of an interval mentioned in Section 3.3.1, if it is desired to only include specific channels in the power spectrum.

#### 3.3.6 Plot

As can be seen in Figure 2A, the majority of the screen is occupied by the plot which dynamically adapts its size based on RV’s window size to fill as much space as possible. All selected channels are spread across the vertical axis with an offset of 40 μV between them by default, unless specified otherwise in the visualization settings. Time, on the horizontal axis, is displayed in seconds. If the view-slider is enabled, the first ten seconds of the recording (or segment) are displayed across all selected channels, by default. Otherwise, the whole recording (or segment) is shown.

The EEG traces are plotted in black, with the exception of bad channels, whose traces are shown in gray. The user can hover over any given point in the plot in order to display the time (in seconds) and amplitude (in μV if no custom scaling was used) values, rounded to three decimals, of the trace under the mouse. When deep-learning predictions are activated in the “Preprocessing” pop-up window (Figure 1), they are plotted below the EEG traces. This will be explained in more detail in Section 3.5. Figure 6A shows an example plot, where the integrated deep-learning model is activated, and several artifacts are annotated.

**Figure 6:**
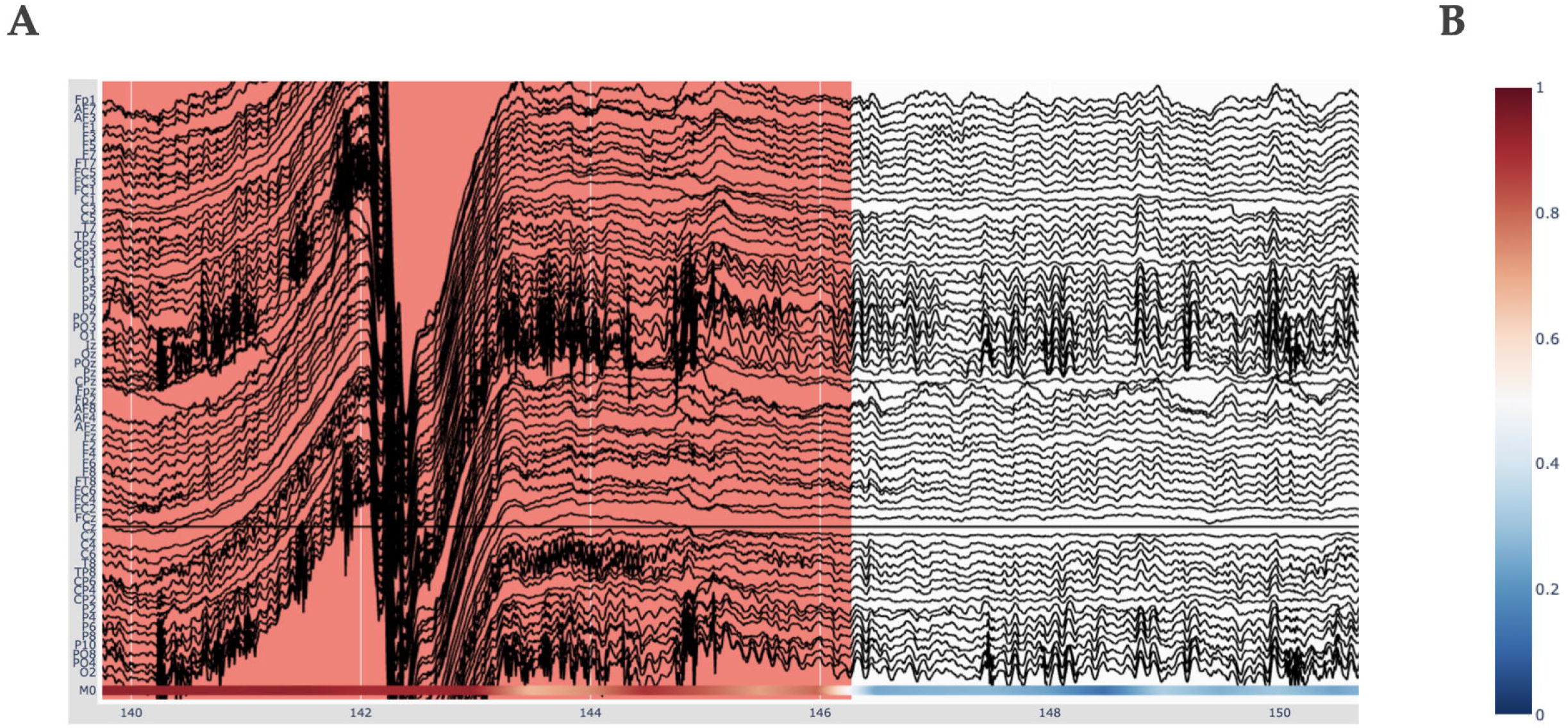
Visualizing deep-learning predictions of artifacts in RV. (A) Example timeframe of an EEG file displayed in RV. An annotated artifact is illustrated here (in red) and the respective deep-learning model predictions at the bottom of the plot, in accordance with the colorbar seen in B. (B) The color indicates predicted probability of an artifact. Red (1) corresponds to high probability and blue (0) to low probability.

### 3.4 Bad-Channel Marking

Bad channels are EEG electrodes which did not record actual brain activity, for example because they were disconnected from the patient’s scalp, and, as such, cannot be used for analysis and interpretation. In RV, we offer two possibilities to mark bad channels: The manual way of inspecting the EEG data first and then marking the bad channels through the GUI, and the automatic method selectable in the “Preprocessing” pop-up window (Figure 1). Bad channels marked in RV can be saved as a list of the respective channel names in a .txt file (see Section 3.7).

#### 3.4.1 Manual Bad-Channel Marking

As mentioned in Section 3.3.2, the user can mark channels as “bad” by clicking on them while the button (#4) in the taskbar (Figure 2B – button 4) is activated. Clicking an already marked bad channel will remove the marking. Additionally, channels can be marked as bad in the respective dropdown-menu in the “Preprocessing” pop-up window (Figure 1), where channels will be selected if the loaded file already contains marked bad channels. In the plot, bad channels will be highlighted in gray. Furthermore, once at least one bad channel has been marked, the “Hide/show bad channels” button (Figure 2A) is enabled. This button allows for bad channels to be hidden from the plot, as explained in Section 3.3.3.

#### 3.4.2 Automatic Bad-Channel Detection

Automatic bad-channel detection is performed using the RANSAC method of the PREP pipeline (Bigdely-Shamlo et al., 2015) implemented in the Autoreject library (Jas et al., 2017), and can be activated in the “Preprocessing” pop-up window (Figure 1). When this method is used, the EEG recording is first cut into 4-second segments, in accordance with the default in the PREP pipeline. Then, 25% of the total number of channels are selected. These selected channels are used to interpolate all other channels. This process is repeated 50 times to generate 50 interpolated time series per channel. Next, for each channel, the correlation of the median of these 50 time series with the corresponding, actual channel is calculated. If this correlation is lower than 75% for more than 40% of the segments of a channel, the respective channel is marked as “bad”. Then, it will automatically be selected in the bad-channel dropdown-menu in the “Preprocessing” pop-up window (Figure 1) and highlighted in gray in the plot. The user can then further adjust the automatically selected channels using the methods outlined above (Section 3.4.1). Note that, for future versions of RV, we are planning to add more automatic bad-channel detection methods.

### 3.5 Deep-Learning Model Predictions

There are two possibilities for visualizing deep-learning model predictions within RV. The first option is to load predictions from a file; the second option is to run the integrated deep-learning model on the currently loaded EEG recording in RV. Once loaded, predictions can be used to automatically annotate the data. Note that RV can also be used without the deep-learning model.

#### 3.5.1 Loading Predictions from Files

The first option for displaying predictions in RV is for the user to run the deep-learning model of choice over each EEG recording individually with their own scripts outside of RV, saving the predictions (scaled between zero and one) in separate files, and then loading them in RV for each recording. Note that there must be one prediction per timepoint in the raw recording. RV assumes that the loaded predictions have the same sampling frequency, and hence align with, the loaded raw EEG data (i.e., without the downsampling applied). If this was not the case, the predictions would not be scaled correctly across the time-axis and RV will display an error message. Note that if the deep-learning model is integrated into RV and run accordingly (see following Section 3.5.2), predictions are internally returned along with their corresponding sampling frequency, allowing for correct scaling along the time-axis.

Loading prediction files is very similar to loading EEG data: After generating the model-prediction files, they must be placed in the “data” directory. To load these files, the user can either drag-and-drop them into the “Drag-and-drop or click here to select model output” area of the “Preprocessing” pop-up window seen in Figure 1, or click on said area, after which a file selection window will open and allow the user to select the file to load (the selected file will still have to be inside the predefined directories as mentioned in Section 3.1). In contrast to loading an EEG recording, however, multiple prediction files can be loaded, allowing the user to compare several models simultaneously. Currently supported file formats for prediction files are .npy and .txt (where every row represents the prediction for one timepoint in the raw recording). Additionally, .csv files containing previously made annotations can be loaded here. These files must have columns called “onset”, “duration”, and “description”, where each row represents one annotation with these respective values. Annotations made in RV can be saved in this format as described in Section 3.7. When the user wants to load an EEG recording other than the one for which the predictions were loaded, or if the user wants to remove the predictions from the plot, they can use the “Remove model predictions” button underneath the dashed file selection area.

#### 3.5.2 Running the Integrated Deep-Learning Model

Underneath the “Remove model predictions” button in Figure 1, there is an option to run the integrated deep-learning model defined in the “run_model.py” file. The model that is currently included was pre-trained to recognize artifacts (Diachenko et al., 2022). Notably, the model receives the raw EEG data as input from RV. This is done to allow for custom preprocessing of the model’s input, independent of the preprocessing applied to the viewed data. In the case of the integrated model by Diachenko et al. (2022), for example, this is necessary because this model specifically requires time-frequency plots generated from segments of the time-series EEG data as input, as mentioned in Section 2.2. Note that, because the model is sensitive to bad channels, they should be marked before running the model and evaluating the predictions.

The code for the integrated deep-learning model can be found in the “model” directory. If the user wants to implement their own model, there are a few steps to follow: In RV, the model predictions are generated in the “run_model” function within the “run_model.py” file. This function receives as input the preprocessed EEG data, established in the “Preprocessing” pop-up window (Figure 1), and the raw EEG data. Depending on whether the user’s model needs a separate preprocessing routine, either input can be used. The output of the “run_model” function has to consist of a one-dimensional array, where each individual prediction is scaled between zero and one, a list of channel names that the predictions are based on, if applicable (otherwise, a list of all channel names or “None” can be returned), and the sampling rate of the predictions (if “None”, the predictions’ sampling rate is assumed to be equal to the sampling rate of the raw EEG data). The latter is required for RV to correctly match the predictions to the time-axis. Following these requirements for the output, the user can simply replace the code in the “run_model” function to load their own model, preprocess the EEG data passed as input to the function as desired, and feed the data to their model to generate the predictions. For more information on how to implement a custom model, please refer to the docstrings in the “run_model.py” file or the documentation on GitHub.

#### 3.5.3 Predictions Visualization

As mentioned previously, the deep-learning models’ predictions are plotted below the EEG channels (Figure 6A) and are colored according to a colorbar (Figure 6B), where the color changes from blue to white to red for predictions ranging from 0 to 0.5 to 1, respectively. Since the integrated deep-learning model was trained to recognize artifacts, in this case, 0 (*blue*) corresponds to a high likelihood of clean data, while 1 (*red*) indicates a high likelihood for an artifact being present. When hovering with the mouse over the predictions, RV shows the time (in seconds) and the corresponding prediction value. In the legend and on the y-axis, if more than one model output is loaded, the prediction channels will be named “M0”, “M1”, etc. If the user loaded previously generated predictions, they will be displayed in the order of the list shown under “Selected model output” in the “Preprocessing” window (Figure 1). If the user ran the integrated model on the data, it will always be the bottom one.

### 3.6 Annotations

In RV, annotations can be made manually using the GUI, or automatically using the integrated (or loaded) deep-learning model predictions. Annotations are saved in the “annotations” attribute of the saved MNE Raw data-structure, each with its own onset-time, duration, and, because we are primarily focused on artifact annotation for this version of RV, with the description “bad_artifact”. The user can save annotations made in RV in a .csv file (see Section 3.7 for more details). Currently, only annotations across all channels are supported. The user will be able to set the description and color of the annotation in the GUI in a future version of RV.

#### 3.6.1 Manual Annotation

As mentioned in Section 3.3.2, annotations can be made only when the button (#5) in the taskbar (Figure 2B – button 5) is activated, which it is by default upon loading. While this mode is active, the user can select a segment of the plot to annotate by clicking and dragging on the plot. Doing so creates a semi-transparent red box spanning the entire vertical axis in view across the selected time interval. Annotations can be freely adjusted in their position and size when selected: the selected annotation can be moved with click-and-drag, and its corners can be dragged in order to adjust its size. If the user wants to delete the annotation, they first have to select it and then click the button (#6) in the taskbar (Figure 2B – button 6) which deletes the currently selected annotation. Note that, if annotations are overlapping, they will be merged in the underlying data structure and will be displayed merged once the plot is redrawn.

#### 3.6.2 Automatic Annotation

It is possible to let the integrated deep-learning model create annotations automatically according to a customizable threshold specified in the “Preprocessing” pop-up window (see Section 3.2 and Figure 1). When doing so, for every datapoint where the supplied model outputs a prediction higher than the given threshold, an annotation will be drawn. These automatic annotations will be given the same description as the manually created ones (“bad_artifact”).

### 3.7 Saving Annotated Data

As mentioned in Section 3.1, RV always keeps one temporary working save-file, which is updated with every annotation made to the data. This file can be loaded in case RV is accidentally closed before saving. This automatic save-file is called “temp_raw.fif” and is located in the main RV directory. To properly save data, the user has to use the “Save to” button in the menu bar (see Figure 2A), which will open a small pop-up window giving the option to either save the data with a given filename and the .fif file-extension, or to overwrite the loaded file (this is possible only when the loaded file is a .fif file in the “save_files” directory; files in the “data” directory cannot be overwritten, as explained in Section 3.1). RV relies on the saving functionality of MNE, which is why we currently only support saving to the native MNE .fif format. For future versions of RV, we will investigate possibilities to export to other EEG-file formats, e.g., .edf and .set (which can currently only be loaded into RV). Once files are saved in the “save-files” directory, they can also be loaded through the same steps as outlined in Section 3.1. If a save-file already exists with the same name, it will be saved with an increasing integer added to the end of the name, unless the user chooses to overwrite the loaded save-file. When choosing to overwrite the loaded save-file, an additional confirmation pop-up window opens.

As mentioned in Section 3.2, the save-file will contain the preprocessed data after applying bandpass filters and setting of the custom reference. This is because the bandpass filter settings can influence the timing of artifacts and we want to avoid that some artifacts are filtered away during viewing but not when subsequently performing further analysis. Visualization settings, such as downsampling, will not be included as these are exclusively used for plotting.

In addition to saving the entire annotated EEG data, there are two buttons to separately save the annotations and bad channels, respectively. The annotations are saved in a .csv file, where each row represents an annotation and there are columns for the onset, the duration, and the description of each annotation (following the same structure MNE uses, as described in Section 3.6). These annotation save-files can subsequently be loaded in RV as external deep-learning predictions (as described in Section 3.5.1), allowing to compare annotations made by multiple deep-learning models or human annotators. However, the user must ensure to keep the preprocessing options constant when loading these annotations due to the potential temporal shifting of artifacts mentioned above. The bad channels are simply saved in a .txt file as a list of the bad channel names. Both files are named according to the full EEG-save files described above, with the addition of “_annotations” and “_bad_channels”, respectively, right before their respective file extension (.csv and .txt).

### 3.8 Integration into EEG Toolboxes

Since RV supports many common EEG-file formats for loading, it can easily be integrated into data-processing pipelines developed with different EEG toolboxes. Aside from loading data in RV and running it through the integrated preprocessing pipeline (Figure 4), the user can preprocess their data in a script outside of RV before loading it or pass it to the “run_viewer” function as a MNE Raw data-structure. The latter allows the user to circumvent the restriction of only loading and saving data in the “data” and “save_files” directories, respectively (for more details on the “run_viewer” function, please refer to the documentation on GitHub). Then, because preprocessing has been done externally, all the preprocessing steps (except the visualization settings) within RV can be skipped (by not changing the preprocessing parameters), and the data can be visualized and annotated immediately. After annotating the data, the user can use the files saved with RV with their custom pipeline afterwards. Importantly, the ability to save the annotations in a separate .csv file also allows for RV to be integrated into MATLAB-based pipelines, even though the entire annotated EEG data can only be saved in .fif files at the moment (see Section 3.7): The data could be preprocessed and saved in a .set file in MATLAB, loaded and annotated in RV, and finally the saved annotations can be loaded back into MATLAB.

## 4 Discussion

We developed an open-source EEG viewer that facilitates EEG annotation with the optional support of deep-learning model predictions. RV offers common EEG viewer functionalities with an emphasis on tools surrounding artifact annotation. The integration of deep-learning model predictions into RV facilitates an efficient decision-support system that can be used in research and clinical settings. An important advantage of RV is that it can be easily integrated into pipelines developed with other EEG toolboxes. Moreover, because RV is programmed in Python, and it gets served in the user’s web browser, RV can be run on any operating system.

The main limitation of RV lies in its performance if neither downsampling nor segmentation is used, as plotting the entire signal can burden memory significantly and result in a long initial loading time and slow-responding GUI. Therefore, for EEG recordings with a high number of channels, or for long EEG recordings, segmenting and downsampling the data (resulting in smaller amounts of data being loaded into memory and plotted at one given time) become indispensable. However, the user can use these settings to adjust the trade-off between RV’s computational cost and the amount of data loaded at one time, which allows RV to be run on a wide range of hardware. Another limitation is that the current version of RV was developed for binary classification (artifact or clean data) and hence does not yet support annotations with customizable labels and colors. Similarly, RV cannot handle multi-class deep-learning predictions yet. However, this will be revised in a future version of RV.

This paper marks the beginning of Robin’s Viewer (Version 1.1). Many additional features and improvements are currently in development for future versions. The main working points are the option to add labels to annotations to support multi-class deep-learning predictions and annotations (e.g., for sleep staging), the possibility to save annotated EEG data in a MATLAB-readable format to better facilitate the integration of RV into MATLAB-based EEG toolboxes, as well as functionality to interact with the RV-GUI using keyboard keys and combinations of keys. Once the implemented deep-learning model is accurate enough, we will also consider adding an option where the user is presented only with segments for which the model’s confidence stays within a certain interval, e.g., predictions between 0.3 and 0.7. This would speed up the annotation process further. Additionally, we want to further reduce the computational cost of RV and improve its speed. In this regard, we are investigating a more sophisticated preloading technique where the data outside of the currently plotted timeframe is loaded on the fly as the user is using the view-slider to scroll through the recording.

Finally, we invite other Python programmers, deep-learning researchers, EEG researchers and clinicians alike to collaborate and build on RV, according to their requirements, or to make feature requests on GitHub at https://github.com/RobinWeiler/RV.git.

## 6 Author Contributions

RW, MD, EJ-M, AEA, PB, and KL-H conceived design and functional requirements. RW wrote the paper; MD, EJ-M, AEA, and KL-H revised and edited the paper. RW wrote the code. MD provided the deep-learning model trained to recognize artifacts (under supervision of PB and KL-H) and code to run it included in RV. RW, MD, EJ-M, and AEA tested the software. KL-H provided data.

## 8 Acknowledgments

We thank the students from the courses “Rhythms of the Brain” (2021) and “Human Neurophysiology” (2022) at the Vrije Universiteit Amsterdam, as well as Jennifer Ramautar and Shilpa Anand at the N=You Neurodevelopmental Precision Center, for testing RV and giving feedback. Furthermore, we thank Laureen Weiler for designing the logo of RV.

This manuscript has previously appeared online as a preprint under DOI: <u>10.1101/2022.08.07.503090</u>

## Footnotes

1 https://github.com/plotly/plotly.py

2 https://dash.plotly.com

